# GoFish: A Streamlined Environmental DNA Presence/Absence Assay for Marine Vertebrates

**DOI:** 10.1101/331322

**Authors:** Mark Y. Stoeckle, Mithun Das Mishu, Zachary Charlop-Powers

**Affiliations:** Program for the Human Environment, The Rockefeller University, New York, New York, United States of America; Hunter College, New York, NY, United States of America; Laboratory of Genetically Encoded Small Molecules, The Rockefeller University, New York, NY, United States of America

**Author notes:** Corresponding author [MYS].

## Abstract

Here we describe GoFish, a streamlined environmental DNA (eDNA) presence/absence assay. The assay amplifies a 12S segment with broad-range vertebrate primers, followed by nested PCR with M13-tailed, species-specific primers. Sanger sequencing confirms positives detected by gel electrophoresis. We first obtained 12S sequences from 77 fish specimens representing 36 northwestern Atlantic taxa not well documented in GenBank. Using the newly obtained and published 12S records, we designed GoFish assays for 11 bony fish species common in the lower Hudson River estuary and tested seasonal abundance and habitat preference at two sites. Additional assays detected nine cartilaginous fish species and a marine mammal, bottlenose dolphin, in southern New York Bight. GoFish sensitivity was equivalent to Illumina MiSeq metabarcoding. Unlike quantitative PCR (qPCR), GoFish does not require tissues of target and related species for assay development and a basic thermal cycler is sufficient. Unlike Illumina metabarcoding, indexing and batching samples are unnecessary and advanced bioinformatics expertise is not needed. The assay can be carried out from water collection to result in three days. The main limitations so far are species with shared target sequences and inconsistent amplification of rarer eDNAs. We think this approach will be a useful addition to current eDNA methods when analyzing presence/absence of known species, when turnaround time is important, and in educational settings.

## Introduction

DNA profiling of ecological communities was first applied to terrestrial microbes [1]. DNA extracted from soil samples—amplified with ribosomal RNA gene primers, cloned, and analyzed by Sanger sequencing—revealed an enormous diversity of uncultured organisms. Whole genome shotgun sequencing provided an alternative culture-independent approach [2]. Combining targeted amplification with high-throughput sequencing, first 454 then Illumina, eliminated cloning and Sanger sequencing, greatly facilitating microbiome study [3-5].

Around the same time, ancient DNA techniques began to be applied to environmental samples, with recovery of 10,000 years-old to 400,000 years-old plant and animal DNA from fecal samples and sediments [6,7]. The earliest reports examining contemporary materials include differentiating human and domestic sources in sewage-contaminated water [8] and recovery of Arctic fox DNA from snow footprints [9]. Taberlet and colleagues were the first to apply an environmental DNA approach to present-day ecology, demonstrating pond water eDNA accurately surveys an invasive frog species [10]. Subsequent work revealed aquatic eDNA detects diverse vertebrates and invertebrates in multiple habitats [11-15]. Aquatic eDNA assays now routinely monitor rare and invasive freshwater species [16-19].

Beginning in 2003, the DNA barcoding initiative firmly demonstrated that most animal species are distinguished by a short stretch of mitochondrial (mt) COI gene [20-22]. This led researchers to assess animal communities by “metabarcoding”, i.e., high-throughput sequencing of mtDNA segments amplified from environmental samples [23-26]. The sequence variability that makes COI an excellent identifier of species hobbles broad-range primer design [27].Primers targeting highly conserved regions in vertebrate 12S or 16S mt genes [28-30] successfully profile aquatic vertebrate communities [31-38]. Multi-gene metabarcoding promises kingdom-wide surveys of eukaryotic diversity [39-43].

Aquatic vertebrate eDNA assays challenge in design and execution. Developing a single-species qPCR test typically necessitates obtaining tissue samples of the target organism and potential confounding species [e.g., 44], and running assays requires a dedicated thermal cycler. High-throughput metabarcoding involves indexing and batching a large number of samples for each sequencing run, and advanced bioinformatics expertise to decode output files.

To facilitate wider use, we aimed for an eDNA assay that did not require tissue samples for validation and could be completed in less than a week. One straightforward technique is species-specific amplification followed by gel electrophoresis and Sanger sequencing, as in early eDNA reports [10]. However, our preliminary experiments generated visible products only in samples with a high number of MiSeq reads, indicating low sensitivity. In addition, multiple bands were frequent, likely to interfere with Sanger sequencing.

Nested PCR is a highly sensitive and specific approach to identifying genetic variants [e.g., 45]. Nested PCR improves detection of earthworm eDNA from soil samples archived for more than 30 years [46] and enables highly sensitive eDNA assays for salmonid fishes and a fresh water mussel [47,48].

We reasoned that nested PCR applied to a metabarcoding target could be a more general approach. Amplification with broad-range metabarcoding primers could provide a “universal” first round, followed by nested PCR with species-specific primers, taking advantage of sequence differences within the amplified segment.

Here we test whether a metabarcoding/nested PCR protocol detects aquatic vertebrate eDNA in time-series water samples from lower Hudson River estuary and southern New York Bight and compare results to those obtained with MiSeq metabarcoding. Because this assay involves querying amplified material one species at a time, we name it after the children’s card game Go Fish in which a player might ask “do you have any Jacks?”

## Results

### New 12S reference sequences

Regional checklist species [49,50] with absent or incomplete GenBank 12S records were flagged. For bony fish, we focused on common species; for the much smaller number of cartilaginous fish we sought specimens from all not in GenBank. Seventy-seven specimens representing 36 target species were obtained from NOAA Northeast Spring Trawl Survey, Monmouth University, fish markets, or beach wrack (Tables S1,S2).

Specimen DNAs were sequenced for a 750 base pair (bp) 12S segment [51] encompassing three commonly used vertebrate eDNA target sites [29,30,32] (Fig 1), and for 648 bp COI barcode region. COI sequences confirmed taxonomic identifications, showing 99.4% to 100% identity to GenBank reference accessions, excepting that of Northern stargazer [*Astroscopus guttatus*], which at the time of this study had no GenBank COI records (Tables S1,S2).

**Fig 1.**
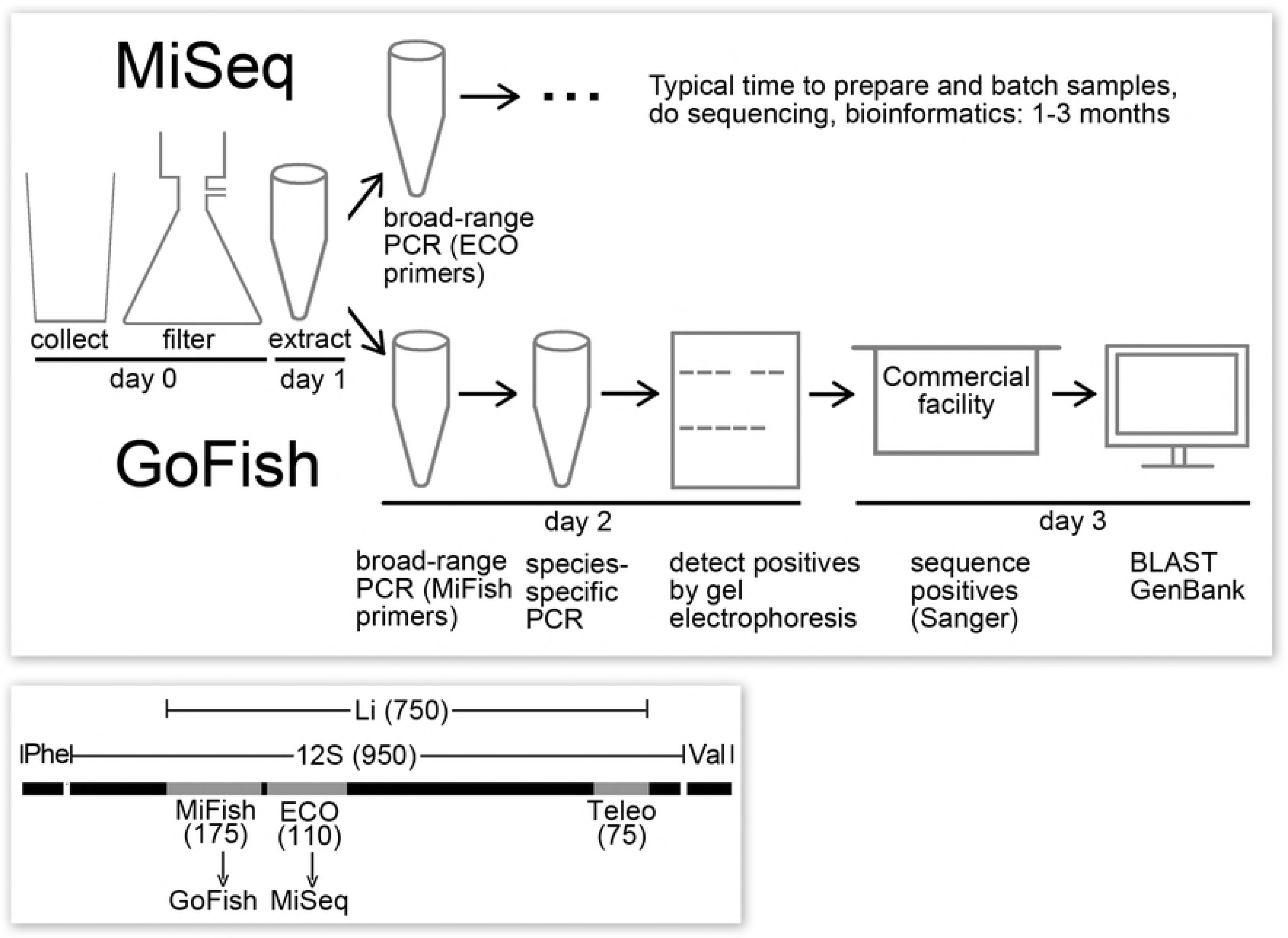
GoFish eDNA assay. Top, schematic of GoFish and MiSeq metabarcoding protocols. Bottom, diagram of 12S and flanking tRNA genes, with locations and sizes of vertebrate metabarcoding targets (MiFish, ECO, Teleo) and the Li segment sequenced from reference specimens as indicated.

### GoFish nested PCR assay

Of three commonly used vertebrate 12S metabarcoding targets (Fig 1), the MiFish segment is longer than the other two and has hypervariable regions near the ends, features facilitating species-specific nested PCR. In addition, by targeting a different segment than what our laboratory uses for MiSeq metabarcoding (12S ECO V5), we aimed to minimize potential cross-contamination between GoFish and MiSeq assays. First-round PCR for bony fish was done with MiFish primer set [30] (Fig 1, Table 1). The resultant reaction mix, diluted 1:20 in Elution Buffer (10mM Tris pH 8.3, Qiagen) served as input DNA for species-specific PCRs.

GoFish primers were designed for 11 fish species that together account for most (92%) lower Hudson River estuary fish eDNA reads (Table 2) [52]. The nested primers generated unambiguous presence/absence bands on gel electrophoresis (Fig 2). In all samples analyzed so far, Sanger sequencing confirmed GoFish amplified only targeted species.

**Fig 2.**
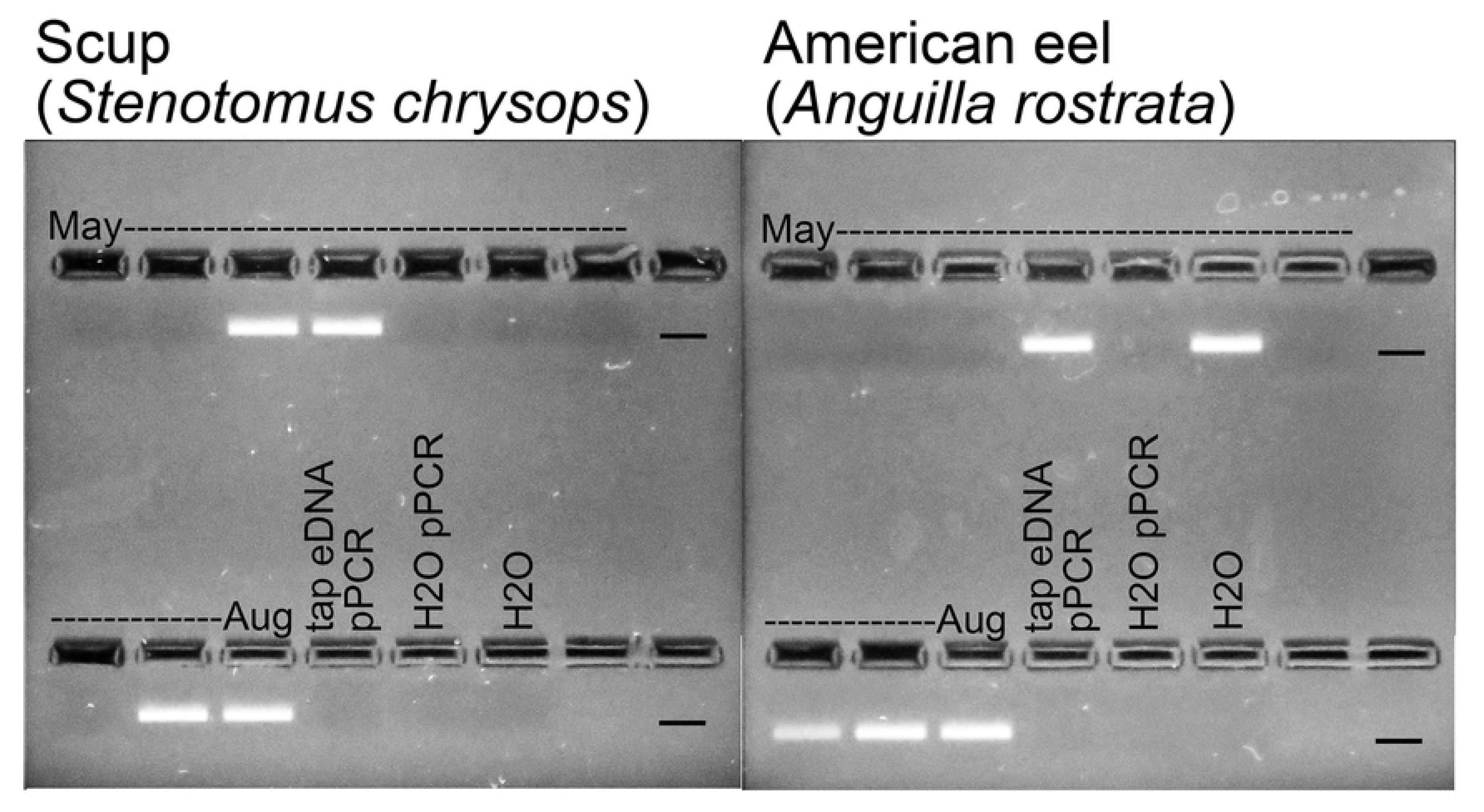
GoFish amplifications visualized on 2.5% agarose gel with SYBER Safe. Lanes bracketed by dates are time series samples; the last three lanes in each panel are negative controls detailed in Materials and Methods. Marker indicates dye front at approximately 150 bp. Gel positives were sent for Sanger sequencing to confirm target species amplification.

**Table 1.**
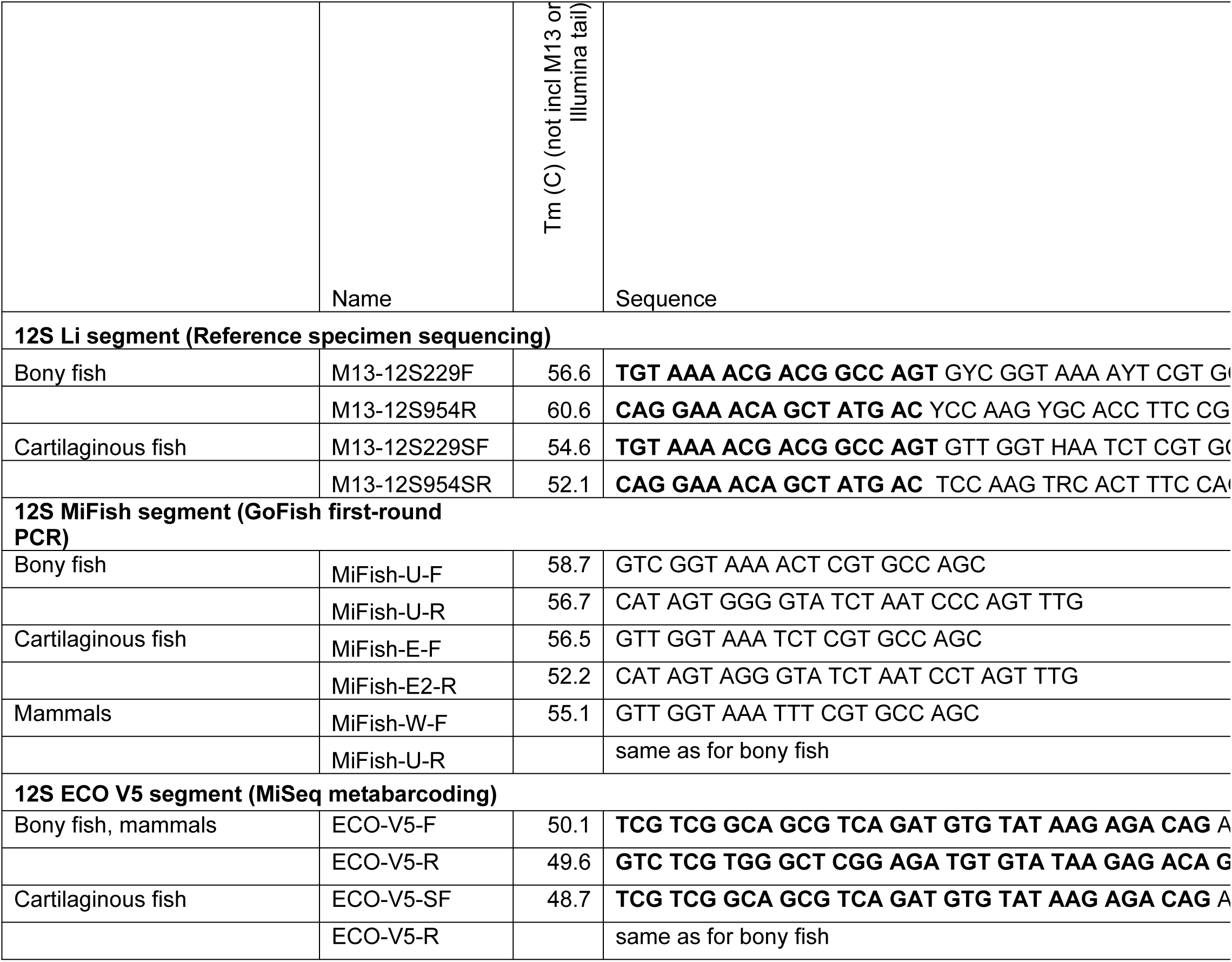
Broad-range vertebrate 12S primers. PCR parameters and expected amplicon sizes are shown. M13 and Illumina tails in Li primers and ECO V5 primers, respectively, are highlighted in bold.

**Table 2.**
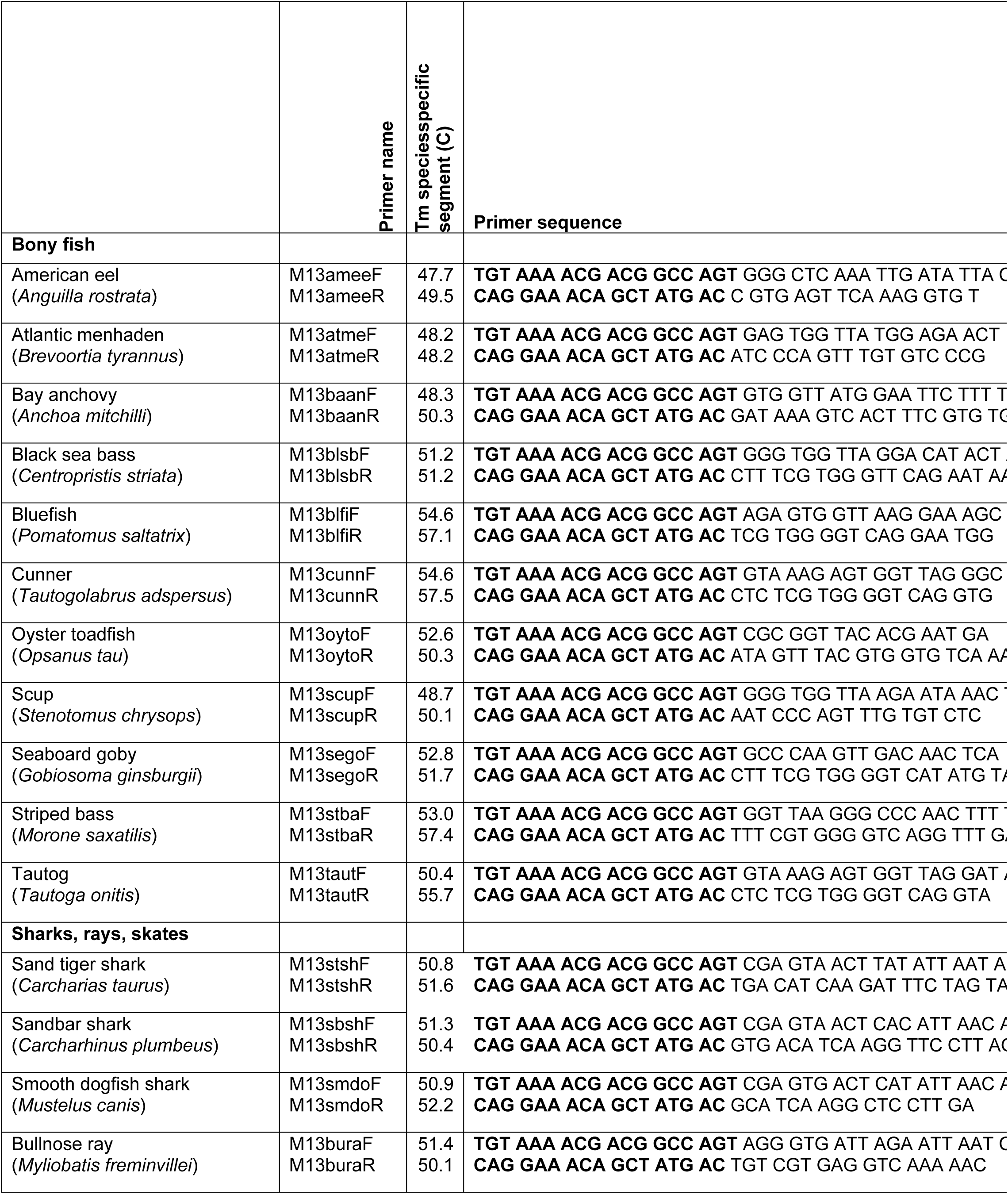
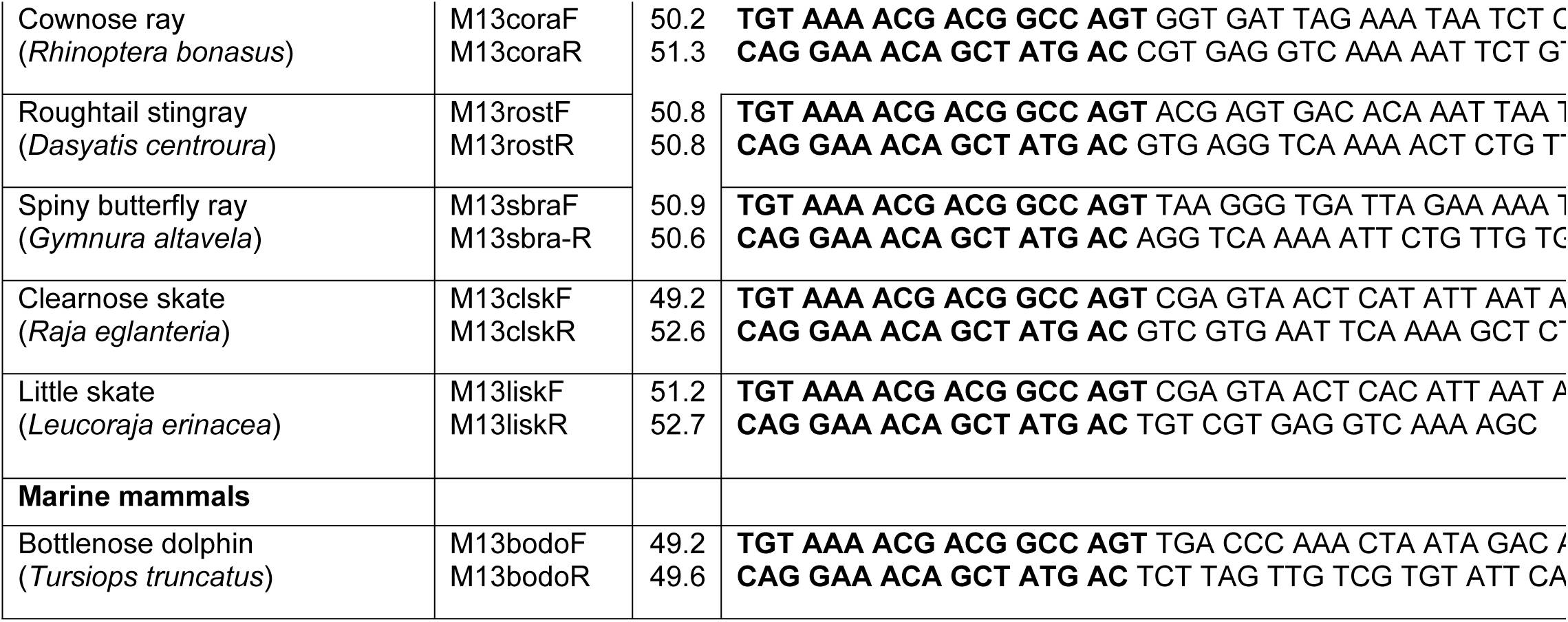
Species-specific GoFish primers. Amplicon sizes, PCR parameters, and target specificity as shown. M13 tails are highlighted in bold.

We applied these GoFish assays to a four-month time series of water samples collected weekly at two contrasting lower Hudson River estuary locations—a high flow, rocky tidal channel on the east side of Manhattan, and a low-flow, sandy bottom site in outer New York harbor (Fig 3]. Species detections increased seasonally at both sites, consistent with historical trawl surveys and a metabarcoding eDNA time series [52]. Despite large tidal flows in the estuary, eDNAs differed by site consistent with habitat preferences, with rocky bottom specialists (cunner, oyster toadfish, seaboard goby) more commonly detected in East River than in outer New York harbor.

**Fig 3.**
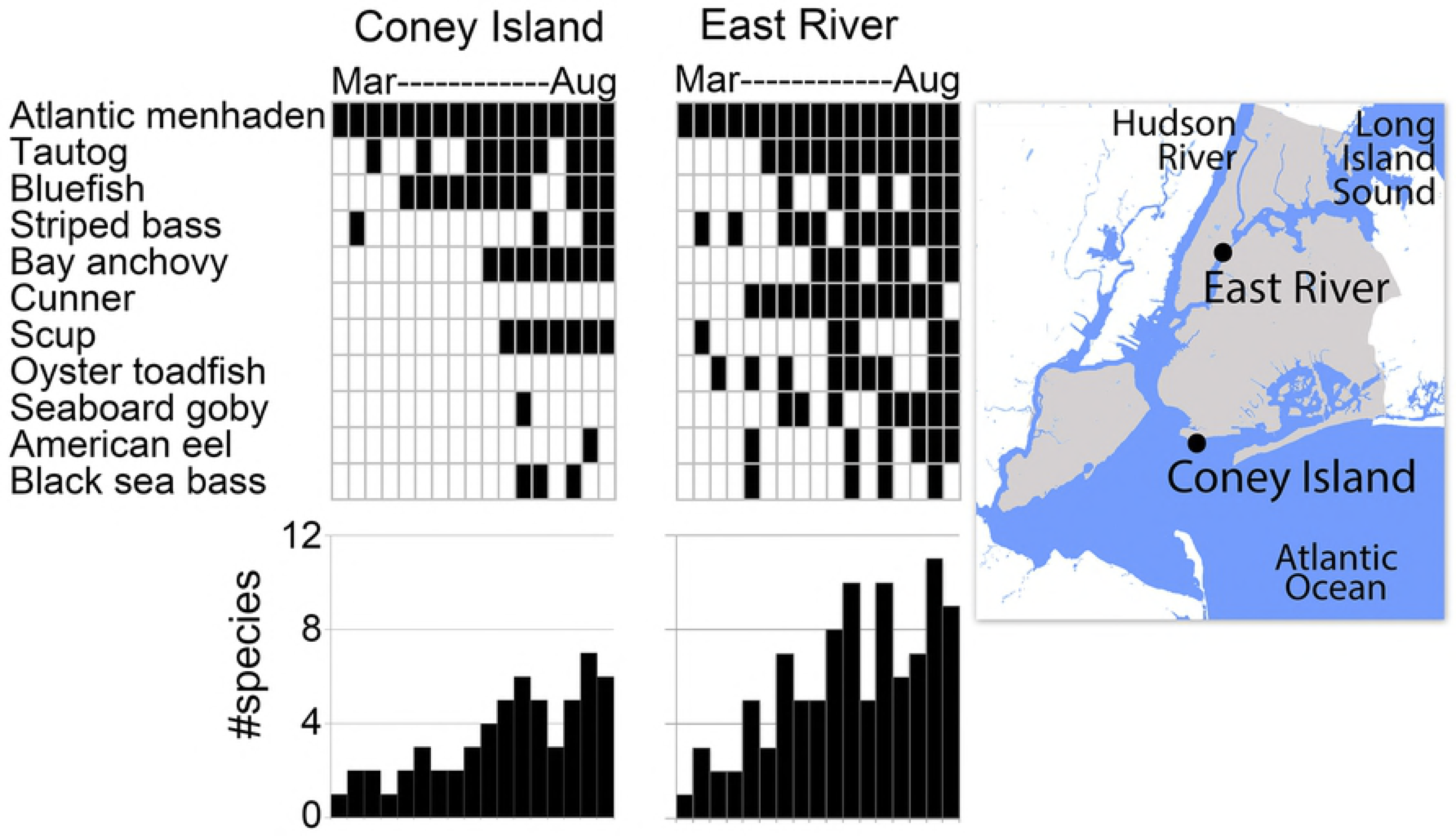
GoFish detections at two lower Hudson estuary locations sampled weekly from March to August 2017. At top, black indicates detections, with species arranged by decreasing number of positives; at bottom, number of species present on each date is shown.

### Cartilaginous fish, marine mammals

We tested this approach on cartilaginous fish and marine mammals, groups relatively understudied by eDNA so far [12,34,36,53-55]. First-round MiFish primers were modified to favor cartilaginous fish or mammals (Table 1). Species-specific GoFish amplifications successfully detected three shark species, four rays, and two skates (Table 2). In a nine-month time series of water samples from southern New Jersey, cartilaginous fish eDNAs peaked in mid-summer, consistent with seasonal migration patterns (Fig 4) [56]. One exception was little skate (*Leucoraja erinacea*), present largely in fall samples, in line with its greater fall and winter abundance [57]. Sanger sequencing confirmed species ID for all gel-positive amplifications, except that clearnose skate (*Raja englanteria*) and little skate primers amplified non-target sequences in some samples. A GoFish assay for bottlenose dolphin (*Tursiops truncatus*) was positive in most summer and fall samples; sequencing verified all (Table 2, Fig 4).

**Fig 4.**
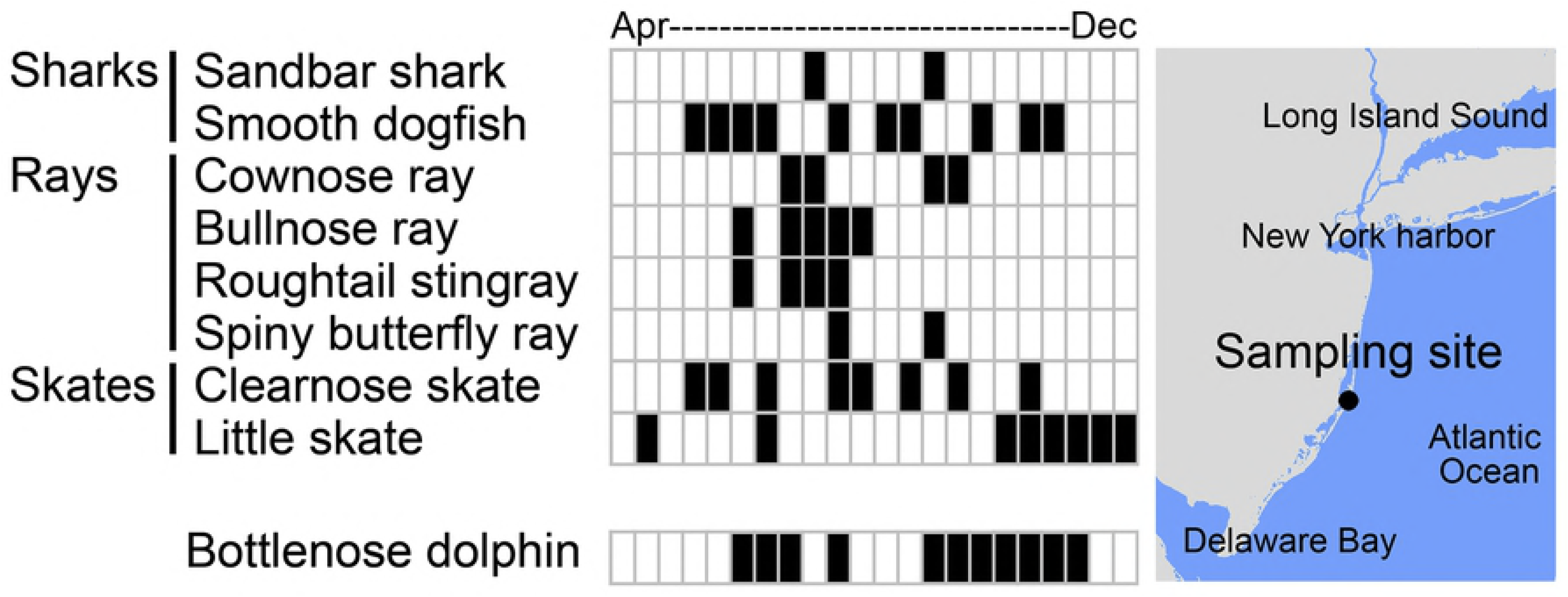
GoFish detections of cartilaginous fish and bottlenose dolphin eDNA. Water samples were collected at one-to two-week intervals from April to December 2017 in southern New York Bight.

### Comparison to Illumina metabarcoding

The New York City time series samples were analyzed by an Illumina MiSeq metabarcoding protocol targeting 12S ECO V5 segment (Fig 1). The apparent sensitivity (method detections/total detections) for both protocols was about 80% (Fig 5). As expected, the proportion of GoFish “drop-outs” differed by MiSeq read number—more abundant eDNAs were amplified more consistently than were rarer eDNAs.

**Fig 5.**
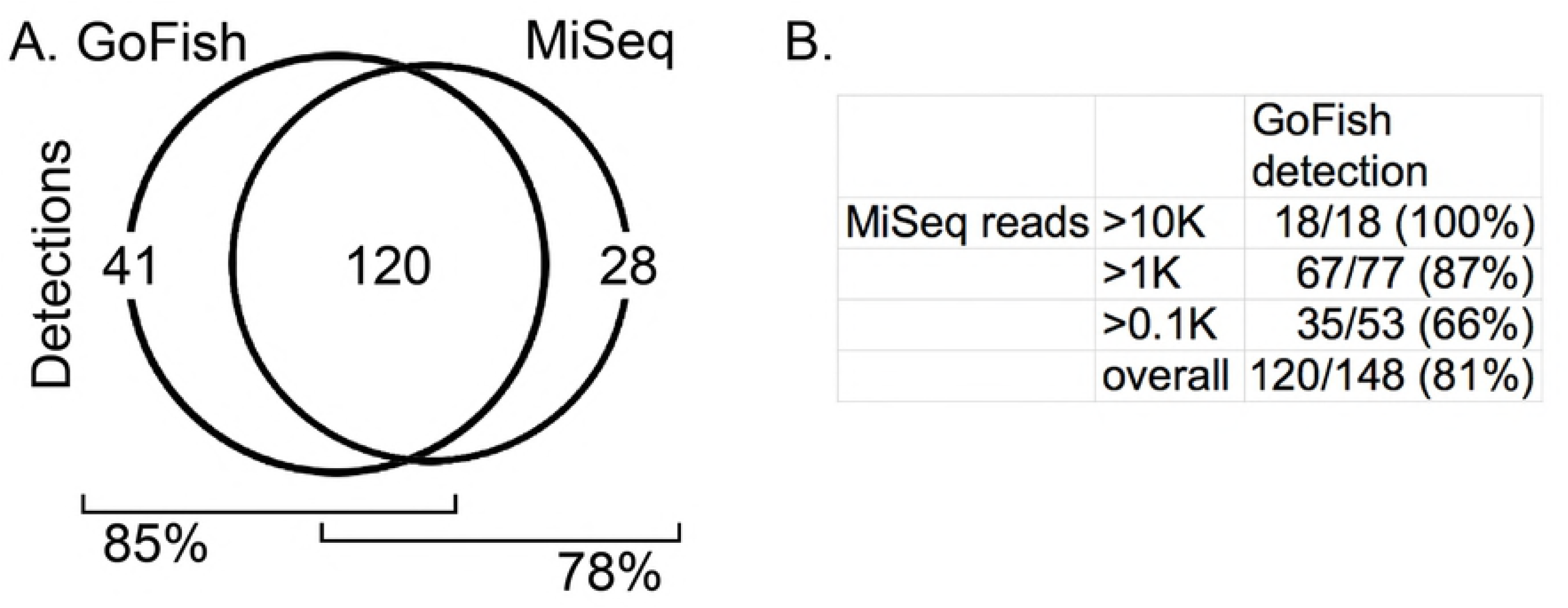
Comparison of GoFish and metabarcoding. A. Method detections for samples and species analyzed in Fig 3. B. GoFish detections stratified by MiSeq read number.

## Discussion

Here we report species-specific nested PCR eDNA assays for 20 marine fish and one marine mammal. The GoFish assay can potentially be adapted to detect any vertebrate with a 12S reference sequence; tissue specimens are not necessary. It can be completed in less than a week with standard molecular biology equipment and interpreted with Sanger sequencing-level bioinformatics. A single broad-range amplification suffices for multiple species-specific assays.

To facilitate primer design we sequenced a 12S fragment, covering three commonly analyzed vertebrate eDNA metabarcoding targets, from 77 specimens representing 36 local species, boosting GenBank 12S coverage to 95% of lower Hudson River estuary checklist species [52].

The main limitations so far are species lacking sequence differences in the target segment (assays require species-specific substitutions in both primer sites and in amplified segment) and inconsistent amplification of rarer eDNAs, shortcomings shared with metabarcoding. GoFish can be viewed as an alternate form of metabarcoding, one that uses species-specific amplification rather than high-throughput sequencing to analyze the pool of PCR products generated by broad-range primers.

Several pairs or sets of local species have insufficient MiFish segment sequence differences for GoFish assays. Of note, these include *Alosa* herrings, of commercial and conservation interest: alewife (*A. pseudoharengus*), American shad (*A. sapidissima*), blueback herring (*A. aestivalis*), and hickory shad (*A. mediocris*). It may be possible to design nested PCR assays for *Alosa* herrings by targeting a different mtDNA region.

GoFish sensitivity was equivalent to that of MiSeq metabarcoding, with drop-outs in both assays (Fig 5). As expected, GoFish drop-outs were mostly those with lower MiSeq read numbers, and presumably represent rarer eDNAs. Inconsistent amplification of low abundance DNAs was recognized as a hazard early on [10]. If desired, replicate amplification or other PCR enhancement strategies [58] could be applied to GoFish. False-negatives may be inherent to broad-range primers, which “rarely detect lineages accounting for less than 0.05% of the total read count, even after 15 PCR replicates” [59] [also 40]. Looked at more generally, all ecological survey methods generate false-negatives; site occupancy modeling can help infer true presence/absence [60-63].

We assumed that sequences matching regional species indicated the presence of that species. This could lead to overlooking extralimital occurrences of taxa with shared sequences. For instance, the locally abundant Atlantic menhaden (*Brevoortia tyrannus*) shares GoFish target sequences with Gulf menhaden (*B. patronus*), found in Gulf of Mexico.

Although likely impractical to apply GoFish to the hundreds of fish species typically resident in any given marine region, it may be possible to characterize communities by targeting the smaller number of species that account for the majority of biomass (e.g., Fig 3).]

The non-labor costs for a GoFish assay were about $15 per sample for one species, and $8 per sample per additional species. One hard to quantify advantage is constrained cross-contamination. Because it is cost-effective to test small sets of samples, a GoFish assay puts fewer results at risk than does high-throughput sequencing. This feature could be particularly valuable in educational settings with less expert performers, instead of putting “all your eggs in one basket” in a MiSeq run.

eDNA promises to help better understand and appreciate ocean life. We think GoFish will be a useful addition to eDNA tools when species of interest are known and relatively few in number, when turnaround time is important, and in educational settings.

## Materials and Methods

The GoFish protocol was designed for persons familiar with basic molecular biology techniques and access to essential molecular biology laboratory equipment. To facilitate use, we utilized commercial kits and open source software, and standardized PCR and sequencing protocols. Procedures were performed on an open bench following routine molecular biology precautions. Particulars include gloves worn for all laboratory procedures and changed after handling water samples and PCR reactions, filtration equipment scrubbed and rinsed thoroughly after each use with tap water, and pipettors and workspace areas wiped with 10% bleach after use. Unfiltered pipette tips were employed; after each procedure used tips were discarded and collection containers rinsed with 10% bleach. Our aquatic eDNA methods are posted online at protocols.io site (https://dx.doi.org/10.17504/protocols.io.p9gdr3w).

## Standardized GoFish 12S amplification protocol

Limited customization of cycle number and annealing temperature was applied (Tables 1,2). Materials and conditions were as follows: GE Illustra beads in 0.2 ml tubes (8 tube strips); 25 µl reaction volume; 5 µl input DNA; 250 nM each primer; and thermal cycler program of 95 °C for 5 m, (25 or 35) cycles of [95 °C for 20 s, (60 °C or 65 °C) for 20 s, 72 °C for 20 s], and 72 °C for 1 m. Primers were obtained from Integrated DNA Technologies (IDT).

### New 12S, COI reference sequences

DNA was extracted from tissues using PowerSoil kit (MoBio). 12S primer sequences and PCR parameters applied to reference specimen DNAs are shown in Table 1. PCR conditions were otherwise as listed in GoFish protocol above. Amplifications were confirmed by agarose gel electrophoresis with SYBER Safe dye (Thermo Fisher Scientific), and PCR clean-up and bidirectional sequencing with M13 primers were done at GENEWIZ. Consensus sequences were assembled in MEGA, using 4Peaks to assess trace files [64,65]. For COI, COI-3 primer cocktail [66], 35 cycles, and 55 °C annealing were used with GoFish protocol outlined above. Substitute forward primers were employed for specimens that failed to generate high-quality sequences with COI-3 cocktail [hickory shad, American shad (M13alosaCOI-F, 5’-TGT AAA ACG ACG GCC AGT TCA ACT AAT CAT AAA GAT ATT GGT AC-3’); windowpane flounder (M13wiflCOI-F, 5’-TGT AAA ACG ACG GCC AGT CTA CCA ACC ACA AAG ATA TCG G-3’)]. 12S and COI reference sequences (Files S1,S2) are deposited in GenBank (Accession nos. MH377759-MH377835 and MH379020-MH379090, respectively).

### Water collection, filtration, DNA extraction

Water sampling was done under permit from New York City Department of Parks and Recreation at two locations: East River (40.760443, -73.956354), a rocky, high-flow tidal channel on the east side of Manhattan, and Steeplechase Pier, Coney Island (40.569576, -73.983297), a sandy bottom, low-flow location in outer New York Harbor (Fig 3). One-liter surface water samples were collected weekly at both sites from March 31, 2017 to August 3, 2017 (34 samples in total).

With authorization from New Jersey Department of Environmental Protection, surface water samples were collected on a barrier island beach (39.741641, -74.112961) about 70 miles south of New York City and halfway to Cape May, the southern border of New York Bight (Fig 3). 22 one-liter samples were collected at one-week to two-week intervals from April 2, 2017 to December 23, 2017.

Samples were filtered within 1 h of collection or stored at 4 °C for up to 48 h beforehand. Water was poured through a paper coffee filter to exclude large particulate matter and then into a filtration apparatus consisting of a 1000 ml side arm flask attached to wall suction, a frittered glass filter holder (Millipore), and a 47 mm, 0.45 µM pore size nylon filter (Millipore). Filters were folded to cover the retained material and stored in 15 ml tubes at -20° C until DNA extraction. As negative controls, one-liter samples of laboratory tap water were filtered and DNA extracted using the same equipment and procedures as for environmental samples.

DNA was extracted with PowerSoil kit with modifications from manufacturer’s protocol to accommodate the filter [52]. DNA was eluted with 50 µl Buffer 6 and concentration measured using Qubit (Thermo Fisher Scientific). Typical yield was 1 µg to 5 µg DNA per liter water filtered.

No animals were housed or experimented upon as part of this study. No endangered or protected species were collected.

## GoFish assay

### Broad-range 12S amplification

Primer sequences, annealing temperatures, and cycle numbers are shown in Table 1. Different MiFish primer sets targeted bony fish, cartilaginous fish, or marine mammals. Tap water eDNA and reagent-grade water were included as negative controls on all amplification sets. After PCR, 5 µl of reaction mixture were run on a 2% agarose gel with SYBER Safe to assess amplification. Rather than affinity bead purification, we diluted the reaction mix 20-fold in Elution Buffer and used 5 µl for nested amplifications, effecting a 100-fold dilution of first-round reaction products. With this protocol, a single broad-range amplification sufficed for 80 species-specific assays.

### Species-specific primer design, amplification, verification

An alignment of 12S MiFish segment sequences from regional fish species including those obtained in this study, marine mammals, and commonly detected non-marine vertebrates (human, pig, chicken, cow, dog, rat), was generated in MEGA using MUSCLE [67], sorted according to a neighbor-joining tree, exported to Excel, and used to generate a matrix showing differences from the consensus [68]. Primers were selected by eye according to desired criteria: two or more nucleotide mismatches against other species at or near the 3’ end, a T_m_ not including M13 tail of 50.0 °C to 52.0 °C according to IDT website, and diagnostic differences within the amplified segment that confirmed target species detection. G-T or T-G primer-template mismatches were considered relatively permissible and so less useful in conferring specificity [30]. M13 tails enabled a single primer set to sequence all detections and improved 5’ end reads; the latter was particularly helpful given the short amplicons generated by GoFish primers (Table 2).

The standardized amplification protocol described above was employed. Default annealing temperature was 60 °C; if non-target amplification occurred, primers were tested at 65 °C. Three negative controls were included in all runs: the two negative controls from the broad-range PCR, and a reagent-grade water blank. A 5 µl aliquot of each PCR reaction was run on an agarose gel with SYBER Safe (Fig 2); positives were sent to GENEWIZ for cleanup and bidirectional sequencing with M13 primers. Sequences were matched to a local file of 12S reference sequences.

### Metabarcoding

As a comparison, eDNA samples were also analyzed by MiSeq metabarcoding protocol using broad-range primers that target 12S ECO V5 segment as previously described (Fig 1) [52]. Briefly, DNA samples from PowerSoil extraction were further purified with AMPure XP (Beckman Coulter) and resuspended in 50 µl of Elution Buffer. Primer sequences and amplification parameters are given in Table 1. Each sample was amplified with ECO V5 primer set as previously described, which effectively targets bony fish and mammals, and with a modified ECO V5 primer set that favors cartilaginous fish (Table 1). As for MiFish amplifications, tap water eDNA and reagent-grade water negative controls were included in all sets. 5 µl of each reaction were run on a 2% agarose gel with SYBR Safe dye. Some negative controls gave faint bands; with MiSeq, these turned out to be human or domestic animal DNA, commonly observed in eDNA work [69].

PCR products were diluted 1:20 in Elution Buffer and Nextera index primers [Illumina] were added following the standardized amplification protocol with 13 cycles and annealing temperature 55 °C. 5 µl of each reaction were run on a 2% agarose gel with SYBR Safe dye to confirm amplification. Indexed PCR libraries were pooled, treated with AMPure XP, and adjusted to 5.4 ng/µl (30 nM assuming 270 bp amplicon) according to Qubit. Sequencing was done at GENEWIZ on an Illumina MiSeq (2 × 150 bp). 56 experimental and 11 control libraries, plus other samples not reported here, were analyzed in three runs with 86-96 libraries per run. Original fastq files with metadata are deposited in NCBI Sequence Read Archive (NCBI BioProject ID PRJNA358446).

Bioinformatic analysis was performed as previously described [52], using DADA2, which identifies all unique sequences rather than lumping according to threshold criteria [70]. DADA2-generated OTU tables and FASTA files of unique sequences are in Supporting Information (Table S3,Files S3-S5). As in prior report, detections representing less than 0.1% of total reads for that sequence were excluded to minimize misassigned reads. Tap water eDNA and reagent-grade water controls were negative for fish eDNA after filtering.

## Acknowledgments

We thank Jesse Ausubel for encouragement and editorial comment, Jeanne Garbarino, Nica Rabinowitz, Doug Heigl, and Odaelys Walwyn for laboratory space and assistance, John Galbraith, Jakub Kircun, and Keith Dunton for fish specimens, and Victor Shahov and Thomas Poku for trialing protocols.

## Supporting Information

**Table S1. Bony fish specimens analyzed for 12S, COI.**

**Table S2. Cartilaginous fish specimens analyzed for 12S, COI.**

**Table S3. DADA2 OTU tables.**

**File S1. New 12S reference sequences. File S2. New COI reference sequences.**

**File S3. DADA2 FASTA file for MiSeq run jun2017.**

**File S4. DADA2 FASTA file for MiSeq run oct2017.**

**File S5. DADA2 FASTA file for MiSeq run jan2018.**

